# Intrabacterial regulation of a cytotoxic effector by its cognate metaeffector promotes *Legionella pneumophila* virulence

**DOI:** 10.1101/2022.06.12.495833

**Authors:** Deepika Chauhan, Ashley M. Joseph, Stephanie R. Shames

**Affiliations:** Division of Biology, Kansas State University, Manhattan, Kansas 66506 USA

**Author notes:** Department of Diagnostic Medicine and Pathobiology, College of Veterinary Medicine, Kansas State University, Manhattan, Kansas 66506 (Deepika Chauhan). Address correspondence to Stephanie R. Shames.

## Abstract

*Legionella pneumophila* is a natural pathogen of unicellular protozoa that can opportunistically infect macrophages and cause Legionnaires’ Disease. Intracellular replication is driven by hundreds of bacterial effector proteins that are translocated into infected host cells by a Dot/Icm type IV secretion system. *L. pneumophila* effectors are temporally regulated in part by a unique family of translocated regulatory effectors, termed metaeffectors, which bind and modulate the function of a cognate effector in host cells. We discovered that regulation of the cytotoxic effector SidI by its metaeffector, MesI, is critical for *L. pneumophila* virulence in natural and opportunistic hosts. MesI binds and negatively regulates SidI activity *in vitro* but how dysregulation of SidI impairs *L. pneumophila* intracellular replication is unclear. Using a chromosomally-encoded inducible expression system, we discovered that dysregulation of SidI, via loss of MesI, was toxic to *L. pneumophila.* SidI enzymatic activity was required for toxicity since *L. pneumophila* growth was unaffected by induced expression of a catalytically inactive *sidI* allele. We found MesI translocation into host cells was dispensable for intracellular replication and that MesI-deficient bacteria were rapidly degraded within host cells. Together, our data suggest a unique role for intrabacterial effector regulation by a translocated metaeffector in *L. pneumophila* virulence.

**Importance:** *Legionella pneumophila* replicates within phagocytic host cells using hundreds of effector protein virulence factors, which canonically subvert the function of host proteins and pathways. *L. pneumophila* encodes a unique family of translocated effectors called metaeffectors, which bind and regulate the function of a cognate effector in host cells. The metaeffector MesI promotes *L. pneumophila* virulence by regulating the cytotoxic effector SidI; however, the MesI regulatory mechanism is poorly understood. We discovered a unique intrabacterial role for MesI in *L. pneumophila* virulence. When uncoupled from MesI, SidI was toxic to *L. pneumophila in vitro* and triggered robust bacterial degradation in host cells. Importantly, translocation of MesI was dispensable for intracellular replication, demonstrating that intrabacterial regulation of SidI contributes to *L. pneumophila* virulence. These data show a unique and important role for MesI regulation of SidI within the bacterium, which challenges the dogma that effectors function exclusively within host cells.

## Introduction

*Legionella pneumophila* is ubiquitous in freshwater environments where it parasitizes and replicates within free-living protozoa. Co-evolution with environmental phagotrophs have conferred on *L. pneumophila* the ability to replicate within mammalian macrophages, which results in a severe inflammatory pneumonia called Legionnaires’ Disease (1–3). Within host cells, *L. pneumophila* replicates within an endoplasmic reticulum (ER)-derived compartment called the *Legionella*-containing vacuole (LCV), which avoids endocytic maturation and fusion with lysosomes (4, 5). Biogenesis of the LCV is facilitated through the activity of over 300 effector proteins, bacterial virulence factors translocated into host cells through a Dot/Icm type IV secretion system (T4SS) (6). Dot/Icm-translocated effectors are critical for *L. pneumophila* virulence in natural and opportunistic hosts (7); however, the role of most effectors remains poorly understood due primarily to functional redundancy and rarity of virulence phenotypes in laboratory infection models.

Many Gram-negative bacterial pathogens employ effector proteins as part of their virulence strategy. Canonically, effector proteins function by targeting host proteins and subverting cellular pathways to the benefit of the pathogen; however, *L. pneumophila* encodes a unique family of translocated effectors termed metaeffectors, which bind and regulate other *L. pneumophila* effectors within host cells (8, 9). Metaeffectors are a key component of *L. pneumophila* pathogenicity since several are required for intracellular replication but how most metaeffectors promote virulence is poorly understood.

We discovered that regulation of the effector SidI (Lpg2504) by its cognate metaeffector, MesI (Lpg2505), is critical for *L. pneumophila* virulence (10). Like most *L. pneumophila* effectors, loss of SidI has no effect on virulence in laboratory infection models; however, loss-of-function mutation in *mesI* (Δ*mesI*) uniquely attenuates *L. pneumophila* virulence in amoebae, macrophages, and mouse models of Legionnaires’ Disease only when SidI is produced (10). SidI is a cytotoxic glycosyl hydrolase that inhibits eukaryotic mRNA translation (11, 12). MesI binds SidI with high affinity, suppresses its toxicity, and blocks SidI-mediated translation inhibition *in vitro* (12). The requirement for MesI is relieved when *sidI* is deleted (Δ*sidI*); however, a single amino acid substitution that renders SidI non-toxic and catalytically inactive (R453P) is also sufficient to restore replication of MesI-deficient *L. pneumophila* (10). These data have provided key biochemical insights into the MesI regulatory mechanism but how MesI regulation of SidI promotes intracellular replication is still unclear.

In this study, we made the surprising discovery that MesI promotes *L. pneumophila* virulence by regulating SidI intrabacterially. SidI was toxic to *L. pneumophila* when uncoupled from MesI *in vitro*. We found that MesI translocation was dispensable for *L. pneumophila* intracellular replication and MesI-deficient bacteria are rapidly degraded in host cells. These data suggest a unique and important role for interkingdom effector regulation in *L. pneumophila* virulence.

## Results

### Overexpression of *sidI* impairs *L. pneumophila* growth *in vitro*

We discovered that the *L. pneumophila* metaeffector MesI promotes bacterial intracellular replication by regulating its cognate effector SidI (10), but how SidI dysregulation attenuates *L. pneumophila* virulence is unclear. A *L. pneumophila* Δ*mesI* mutant is impaired for intracellular replication but virulence is restored by *sidI* chromosomal deletion (Δ*sidI*Δ*mesI*) or catalytic inactivation (*sidI*_R453P_Δ*mesI*). We initially validated that dysregulation of SidI impairs *L. pneumophila* intracellular replication by plasmid-based genetic complementation of the Δ*sidI* mutation. We infected primary bone marrow-derived macrophages (BMDMs) from *Nlrc4*^-/-^ mice, which are permissive to flagellated *L. pneumophila* (13, 14). Indeed, induced expression of *sidI* from a complementing plasmid severely attenuated *L. pneumophila* replication within BMDMs (**Fig 1A**). Interestingly, we had to induce *sidI* expression at the time of infection since we were unable to culture *L. pneumophila* Δ*sidI*Δ*mesI* (p*sidI*) on inducing media. Inducible plasmid-based genetic complementation is an established technique to fulfill molecular Koch’s postulates and we routinely culture *L. pneumophila* on inducing media (10, 12, 15–17). We hypothesized that overexpression of SidI adversely affects bacterial physiology. Indeed, *L. pneumophila* harboring p*sidI* was unable to replicate in broth under inducing conditions but not in the absence of induction (**Fig 1B**). Impaired growth resulted from SidI enzymatic activity since induced the catalytically inactive *sidI*_R453P_ allele did not affect *L. pneumophila* growth (**Fig 1B**). Our initial interpretation was that dysregulation of SidI does not adversely affect *L. pneumophila* physiology since *L. pneumophila* Δ*mesI* is not attenuated for growth *in vitro* (10). However, *sidI* expression in broth grown bacteria is ∼ 3-fold lower than expression during intracellular replication (18), which may be insufficient to impair *L. pneumophila* growth. Induced expression of an effector gene has not previously been associated with impaired bacterial growth and these data suggest that SidI enzymatic activity is deleterious to *L. pneumophila* when stoichiometrically uncoupled from MesI.

**Figure 1.**
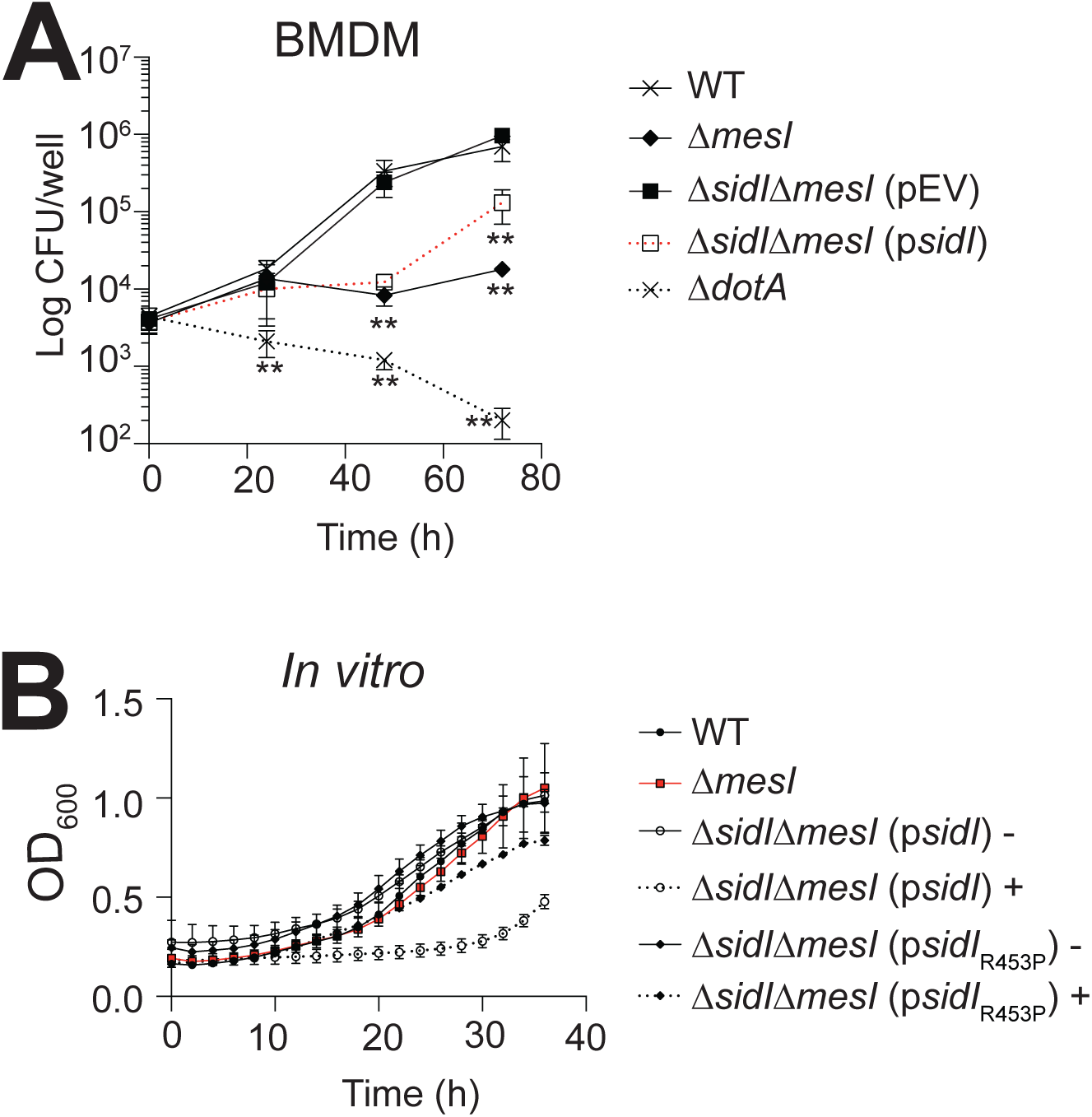
Overexpression of *sidI* attenuates *L. pneumophila* replication. **(A)** Fold replication of *L. pneumophila* strains within *Nlrc4*^-/-^ BMDMs over 72 h. Where indicated, infections were performed in the presence of 1 mM IPTG. Data shown are mean ± s.d. on samples in triplicates and asterisks denote statistical significance by Students’ *t*-test (\*\**P*<0.01). **(B)** Optical density at 600 nm (OD_600_) of *L. pneumophila* strains grown in AYE broth. Where indicated, bacteria were grown in the presence (+) or absence (-) of 1 mM IPTG. Data shown are mean ± s.d. on samples in triplicates and asterisks denote statistical significance by Students’ *t*-test (\*\**P*<0.01). Data shown are representative of at least three independent experiments.

### MesI rescues the SidI-mediated growth defect *in vitro*

MesI suppresses SidI toxicity in yeast (10); thus, we hypothesized that MesI also suppresses SidI toxicity in *L. pneumophila*. To test this hypothesis, we generated *L. pneumophila* strains that enable tightly controlled inducible expression of the *sidI-mesI* locus from its endogenous position in the chromosome. We inserted the *araC-araBAD* (araBAD) promoter and an in-frame *3xflag* epitope tag directly upstream of *sidI* in the chromosome of WT, Δ*mesI*, and *sidI*_R453P_Δ*mesI L. pneumophila* strains (**Fig 2A**). Kinetics and abundance of 3xFLAG-SidI fusion protein production was consistent between strains and only observed under arabinose-inducing conditions (**Fig 2B**). We observed a tightly controlled increase in araBAD-mediated *sidI* expression relative to endogenous levels (**Fig 2C**). The ∼4-fold increase in *sidI* expression from the araBAD promoter is more physiologically relevant than the >10-fold increase in expression from the P*tac* promoter (**Fig 2C**) (18). Thus, we have established *L. pneumophila* strains in which we can control expression of *sidI* and *mesI* in their native stoichiometry and endogenous locus in the chromosome.

**Figure 2.**
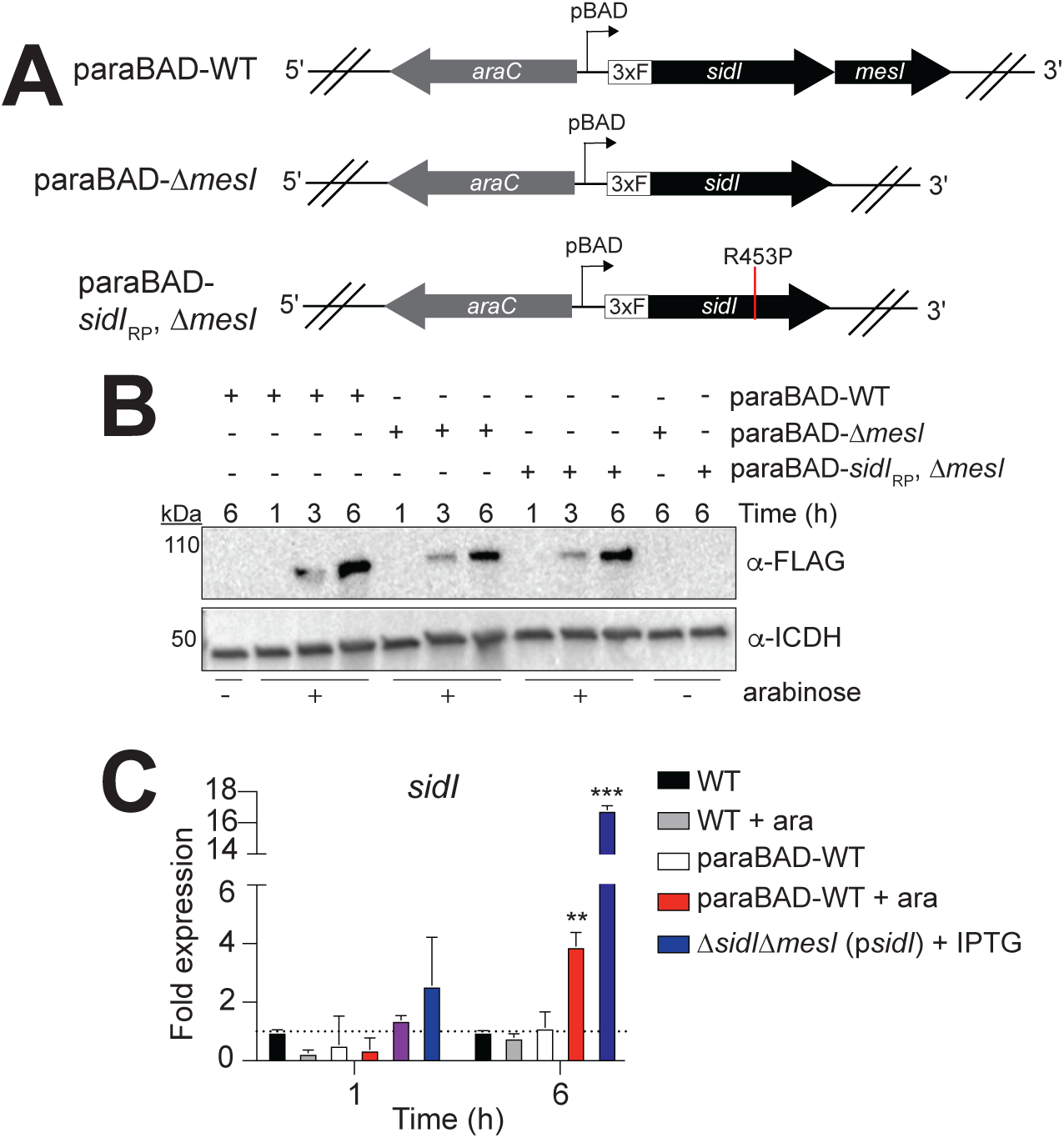
Establishment of a chromosomal arabinose-inducible expression system in *L. pneumophila*. **(A)** Schematic showing paraBAD insertion upstream of the sidI locus in the *L. pneumophila* chromosome. 3XF: In-frame *3xflag* epitope tag. **(B)** Western blot showing 3xFLAG-SidI production in the indicated paraBAD strains in the presence (+) or absence (-) of arabinose [ara; 0.6% (w/v)] after growth in AYE broth for 1, 3, or 6 h. **(C)** Quantitative RT-PCR for expression of *sidI* grown for 1 or 6 h in AYE broth. Fold expression in OD-normalized cultures (*n* = 3) was calculated relative to wild-type *L. pneumophila* at each time point. Asterisks denote statistical significance by Students’ t-test (\*\**P*<0.01) on samples in triplicates. Data are representative of at least two independent experiments.

We leveraged our *L. pneumophila* araBAD strains (**Fig 2A**) to test the hypothesis that stoichiometric production of MesI abrogates SidI-mediated growth attenuation. We quantified replication our *L. pneumophila* araBAD strains under inducing and non-inducing conditions and found that induced expression of *sidI,* but not the inactive *sidI*_R453P_ allele, significantly attenuated bacterial growth only when uncoupled from *mesI* in broth culture (**Fig 3A**) and within BMDMs (**Fig 3B**). Thus, the SidI-mediated growth defect is suppressed by MesI.

**Figure 3.**
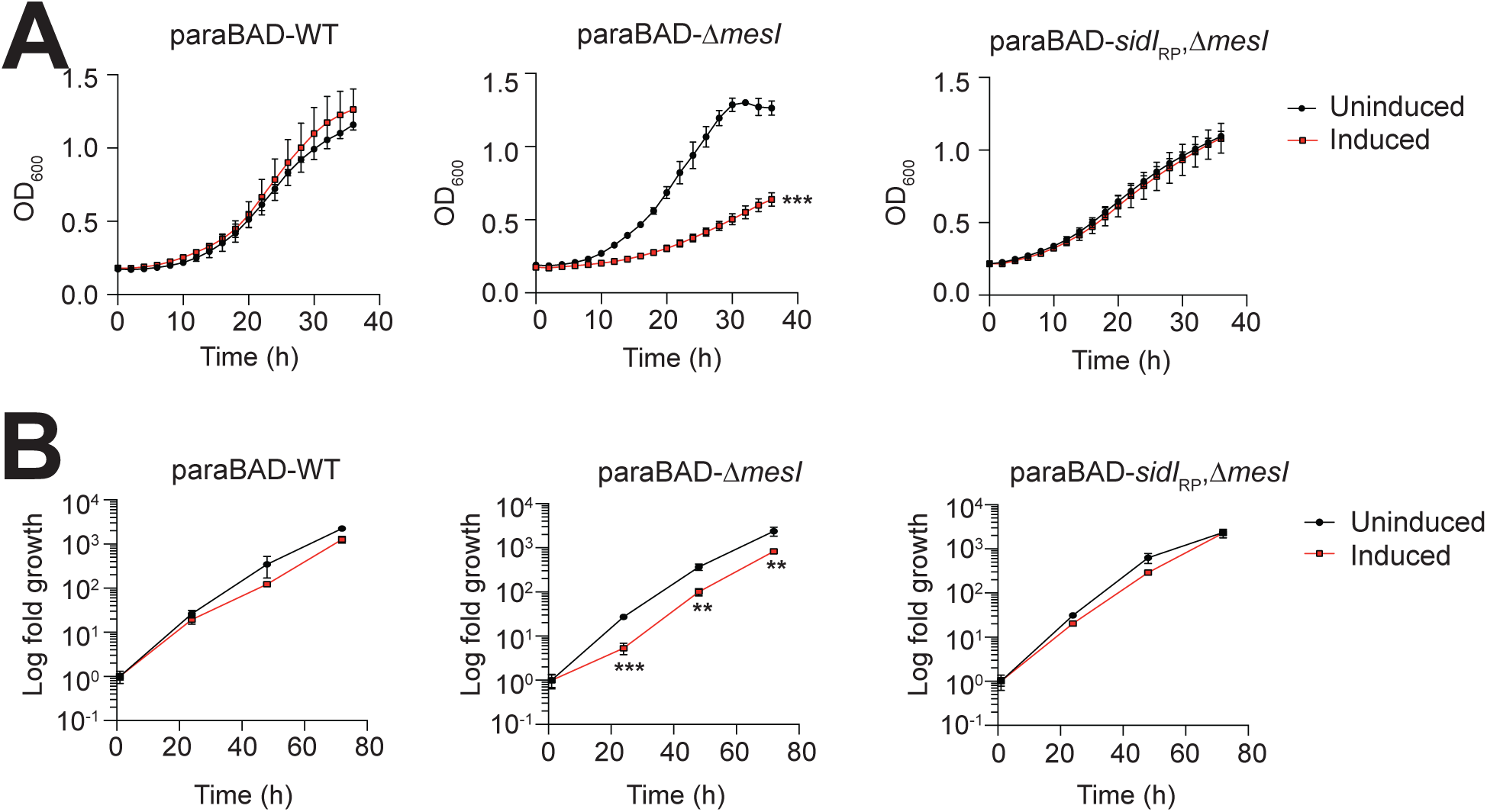
SidI impairs *L. pneumophila* growth when uncoupled from MesI. **(A)** Replication of the *L. pneumophila* paraBAD strains grown in AYE broth *in vitro* under inducing [1% arabinose (w/v)] or noninducing conditions. Data shown are mean ± s.d. OD600 values from samples in triplicates and asterisks denote statistical significance by Students’ t-test (\*\**P*<0.01). **(B)** Fold replication of *L. pneumophila* strains within *Nlrc4*^-/-^ BMDMs under inducing [2% arabinose (w/v)] or noninducing conditions. Data shown are mean ± s.d. on samples in triplicates and asterisks denote statistical significance by Students’ *t*-test (\*\**P*<0.01). Data shown are representative of at least three independent experiments.

### MesI suppresses SidI intrabacterial toxicity

SidI impairs *L. pneumophila* growth, but it is unclear whether SidI is bacteriostatic or bactericidal. SidI is toxic to yeast (11); thus, we tested the hypothesis that SidI is bactericidal when uncoupled from MesI. *L. pneumophila* araBAD strains (**Fig 2A**) were grown for 24 h under arabinose inducing or non-inducing conditions in broth and viability was evaluated using LIVE/DEAD viability staining and confocal microscopy (**Fig 4**). Very few dead bacteria were observed under non-inducing conditions or when induced *sidI* expression was coupled with *mesI* (araBAD-WT) (**Fig 4**). However, significantly more dead bacteria were observed when enzymatically active *sidI* was uncoupled from *mesI* under inducing conditions (**Fig 4**). These data suggest that SidI toxicity is conserved across the kingdoms of life and suppressed by MesI.

**Figure 4.**
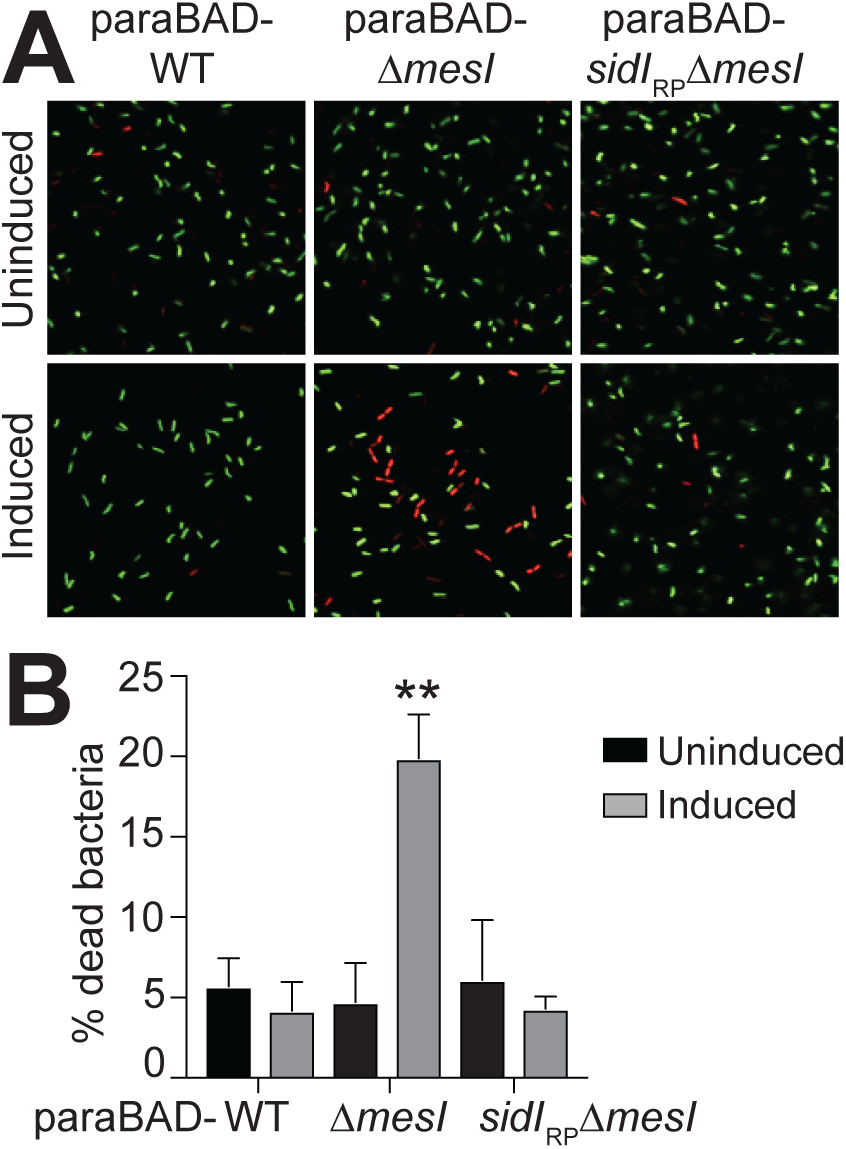
SidI is bactericidal in the absence of MesI. **(A)** Representative confocal micrographs of Live/Dead-stained *L. pneumophila* paraBAD strains grown in AYE broth under inducing [0.6% arabinose (w/v)] or noninducing conditions [SYTO9 (live; green) and propidium iodide (PI; dead; red)]. **(B)** Quantification of dead *L. pneumophila* by fluorescence microscopy following Live/Dead staining. Percent dead bacteria was calculated by dividing PI-stained cells by total cells. Data are mean ± s.d. of 650 cells scored blind and asterisks denote statistical significance by Students’ *t*-test (\*\**P*<0.01). Data are representative of two independent experiments.

### Intrabacterial regulation of SidI by MesI is sufficient for *L. pneumophila* virulence

Based on our observation that SidI is toxic to *L. pneumophila* when uncoupled from MesI, we hypothesized that MesI intrabacterial regulation of SidI is sufficient for *L. pneumophila* virulence. We tested this by evaluating whether an intrabacterially-retained MesI mutant could complement the Δ*mesI* mutation. We generated an allele of *mesI* lacking 10 amino acids from its extreme carboxy terminus; these amino acids are dispensable for SidI binding and contains the putative E-block Dot/Icm translocation signal (19, 20). We cloned this allele into a complementing plasmid downstream of a *3xflag* epitope tag (p*mesI*_Δ10_). We evaluated 3xFLAG-MesIΔ10 translocation using Western blot to visualize its abundance in saponin-soluble or -insoluble lysate fractions from *L. pneumophila*-infected cells, a technique routinely used to evaluate effector secretion, since saponin solubilizes eukaryotic membranes but minimally lyses *L. pneumophila* (21, 22). HEK293 FcγRII cells were infected with antibody-opsonized *L. pneumophila* harboring either p*mesI* or p*mesI*_Δ10_. 3xFLAG-MesIΔ10 was not translocated since it was retained in the insoluble fraction (pellet) and not present in saponin-soluble lysate fractions (**Fig 5A**). As expected, wild-type MesI was secreted into host cells since it was present within saponin-soluble lysate fractions (**Fig 5A**). Isocitrate dehydrogenase (ICDH) and β-actin were used as bacterial lysis and loading controls, respectively. We also confirmed that MesIΔ10 interacts with SidI since it was retained by SidI on beads similarly to full-length MesI (**Fig S1**).

**Figure 5.**
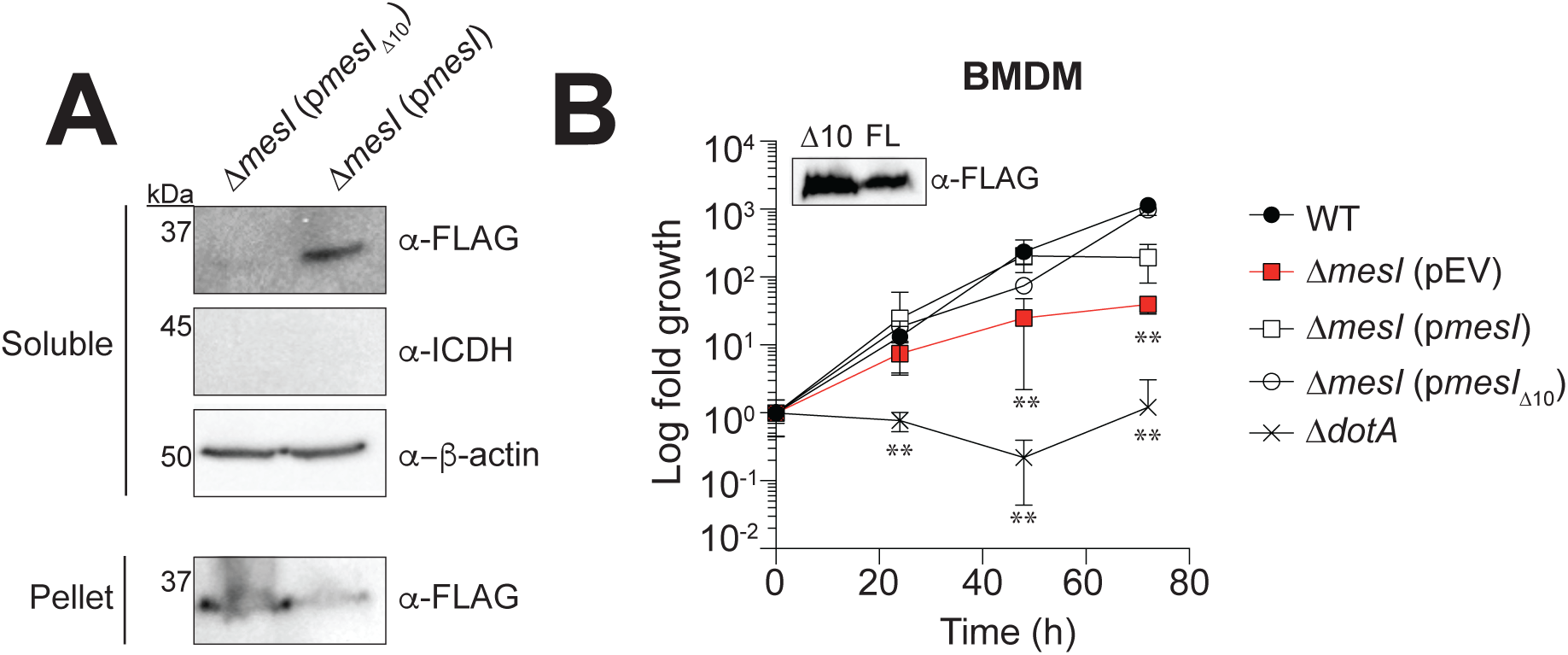
MesI translocation is dispensable for *L. pneumophila* intracellular replication. **(A)** Western blot for 3xFLAG-MesI fusion proteins in saponin-solubilized lysates of HEK293 FcγRII cells infected with antibody-opsonized *L. pneumophila* Δ*mesI* harboring p*mesI* (3xFLAG-MesI) or p*mesI*_Δ10_ (3xFLAG-MesIΔ10). Expression of 3xFLAG-MesI fusions was confirmed by Western blot on saponin-insoluble pellets. ICDH and β-actin were used as controls for bacterial lysis and equal loading, respectively. Data are representative of three independent experiments. **(B)** Replication of *L. pneumophila* strains within *Nlrc4*^-/-^ BMDMs infected at an MOI of 1. Fold growth was calculated by normalizing CFU counts to internalized bacteria at 1 h post-infection. Plasmid expression of *mesI* alleles was induced with 1 mM IPTG and confirmed by Western blot (inset). Asterisks denote statistical significance (\*\**P*<0.05) by Students’ *t*-test on samples in triplicates. Data are representative of three independent experiments.

To evaluate a role for intrabacterial SidI regulation by MesI in *L. pneumophila* virulence, we tested whether the *mesI*_Δ10_ allele genetically complements that Δ*mesI* mutation. We infected BMDMs and quantified replication of wild-type *L. pneumophila* and Δ*mesI* strains harboring p*mesI*, p*mesI*_Δ10_, or empty plasmid vector (pEV). We were able to genetically complement the Δ*mesI* with the *mesI*_Δ10_ allele, demonstrating that MesI translocation into host cells is not necessary for *L. pneumophila* intracellular replication (**Fig 5B**). These data suggest that intrabacterial regulation of SidI by MesI is sufficient for *L. pneumophila* virulence.

### MesI-deficient bacteria are degraded within host cells

We found that MesI suppresses intrabacterial SidI toxicity and that MesI translocation into host cells is dispensable for *L. pneumophila* intracellular replication. Since SidI is also toxic to eukaryotic cells, we initially postulated that the virulence defect associated with loss of MesI results from SidI toxicity in host cells. However, we were unable to detect SidI-mediated cytotoxicity in *L. pneumophila*-infected BMDMs (**Fig S2**) and our new data suggest a potential role for MesI suppression of intrabacterial SidI toxicity in *L. pneumophila*’s virulence strategy.

Dead and dying bacterial are rapidly degraded within host lysosomes; thus, we rationalized that MesI-deficient bacteria would be robustly degraded within host cells. Degraded *L. pneumophila* are easily identified using immunofluorescence microscopy since they have lost their rod shape and appear instead as diffuse puncta when cells are stained with a *Legionella*-specific antibody (23, 24). Thus, we used immunofluorescence microscopy and blinded scoring to quantify degradation of *L. pneumophila* Δ*mesI* within BMDMs. We infected BMDMs and compared kinetics of *L. pneumophila* Δ*mesI* degradation to virulent *L. pneumophila* and *L. pneumophila* Δ*dotA*, which is unable to evade lysosomal degradation (25). After 1 h of infection, most bacteria retained their rod shape and there were no differences between *L. pneumophila* strains; however, at 8 h post-infection, significantly more *L. pneumophila* Δ*mesI* were degraded compared to both wild-type and Δ*dotA* control strains (**Fig 6**). It is unlikely that bacterial degradation was accompanied by host cell death since macrophages harboring degraded bacteria had normal nuclear morphology (**Fig 6**). The robust degradation of *L. pneumophila* Δ*mesI* compared to the avirulent Δ*dotA* strain suggests a mechanism of bacterial attenuation distinct from the inability to subvert endocytic maturation.

**Figure 6.**
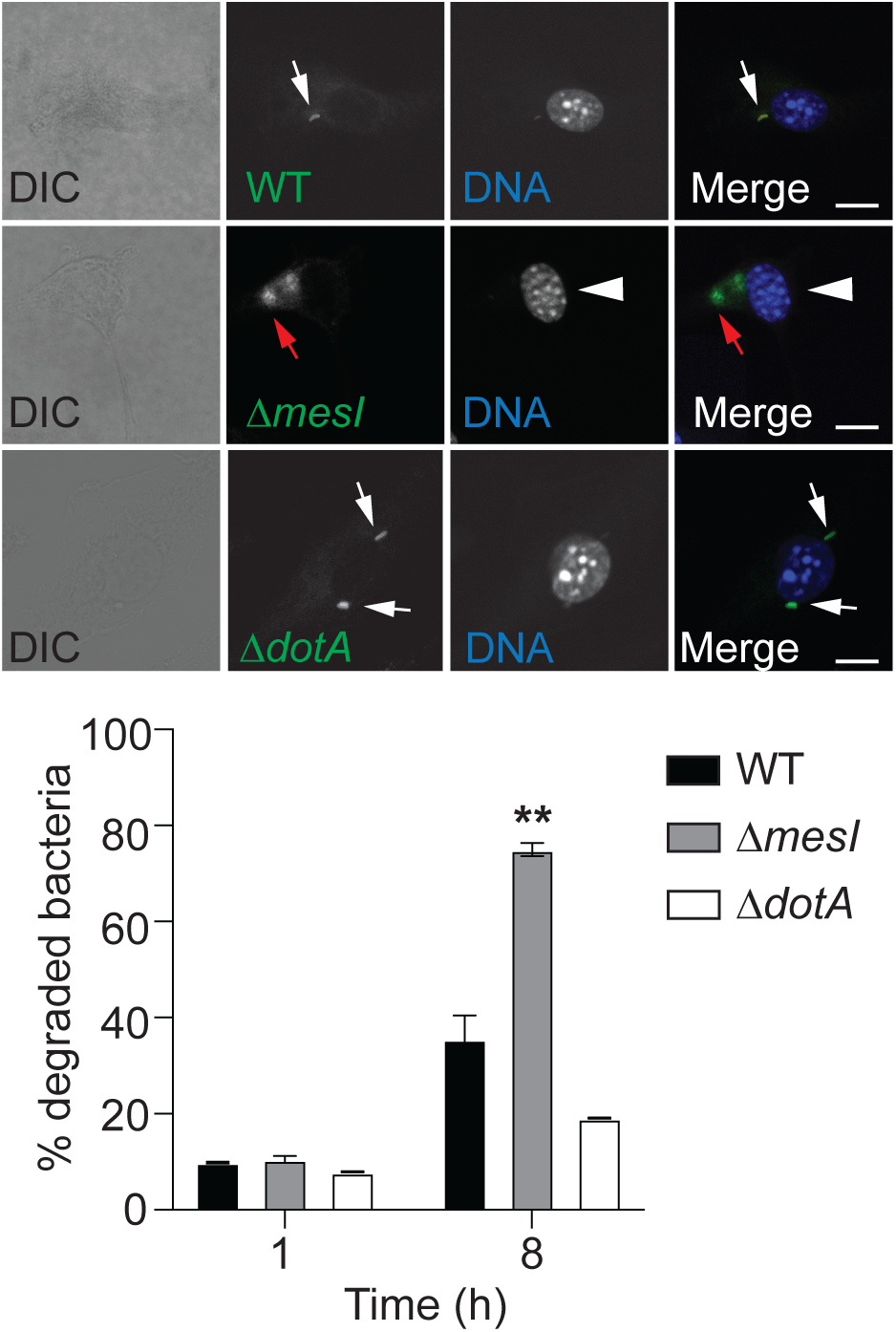
MesI-deficient bacteria are efficiently degraded in host cells. **(Top)** Representative confocal micrographs of *Nlrc4*^-/-^ BMDMs infected for 8 h (MOI of 30) with *L. pneumophila* wildtype (WT), Δ*mesI*, or Δ*dotA*. Intact and degraded bacteria (green) are indicated with white and red arrows, respectively. Nuclei of Δ*mesI*-infected BMDMs are indicated with white arrowhead. Scale bar is 10 µm. **(Bottom)** Quantification of degraded *L. pneumophila* within *Nlrc4*^-/-^ BMDMs infected for 1 or 8 h at an MOI of 30. were infected with *L. pneumophila.* Percent degraded bacteria were enumerated by blinded immunofluorescence scoring of samples in triplicates (*n* = 300). Asterisks denote statistical significance (\*\**P*<0.01) by Students’ *t*-test and data shown are representative of two independent experiments.

## Discussion

This study supports a unique model whereby the *L. pneumophila* metaeffector MesI drives virulence by intrabacterial regulation its cognate effector SidI (**Fig 7**). Several *L. pneumophila* effectors are toxic to eukaryotic cells; however, effector bactericidal activity in *L. pneumophila* has not been previously observed. Furthermore, our data challenge the dogma that metaeffectors promote *L. pneumophila* virulence by functioning exclusively within host cells. SidI and MesI bear resemblance to canonical toxin-antitoxin (TA) and effector-immunity (E-I) pairs (26–28); however, they are distinct since anti-toxin/immunity proteins, by definition, are not secreted(29). This also distinguishes MesI from canonical chaperones, which remain intrabacterial (30). MesI negatively regulates SidI subversion of eukaryotic homeostasis (12) and a role for host cell pathways in the MesI regulatory mechanism cannot be fully excluded (**Fig 7**). However, our data suggest a novel mechanism by which *L. pneumophila* virulence hinges on effector regulation by a metaeffector within the bacterium.

**Figure 7.**
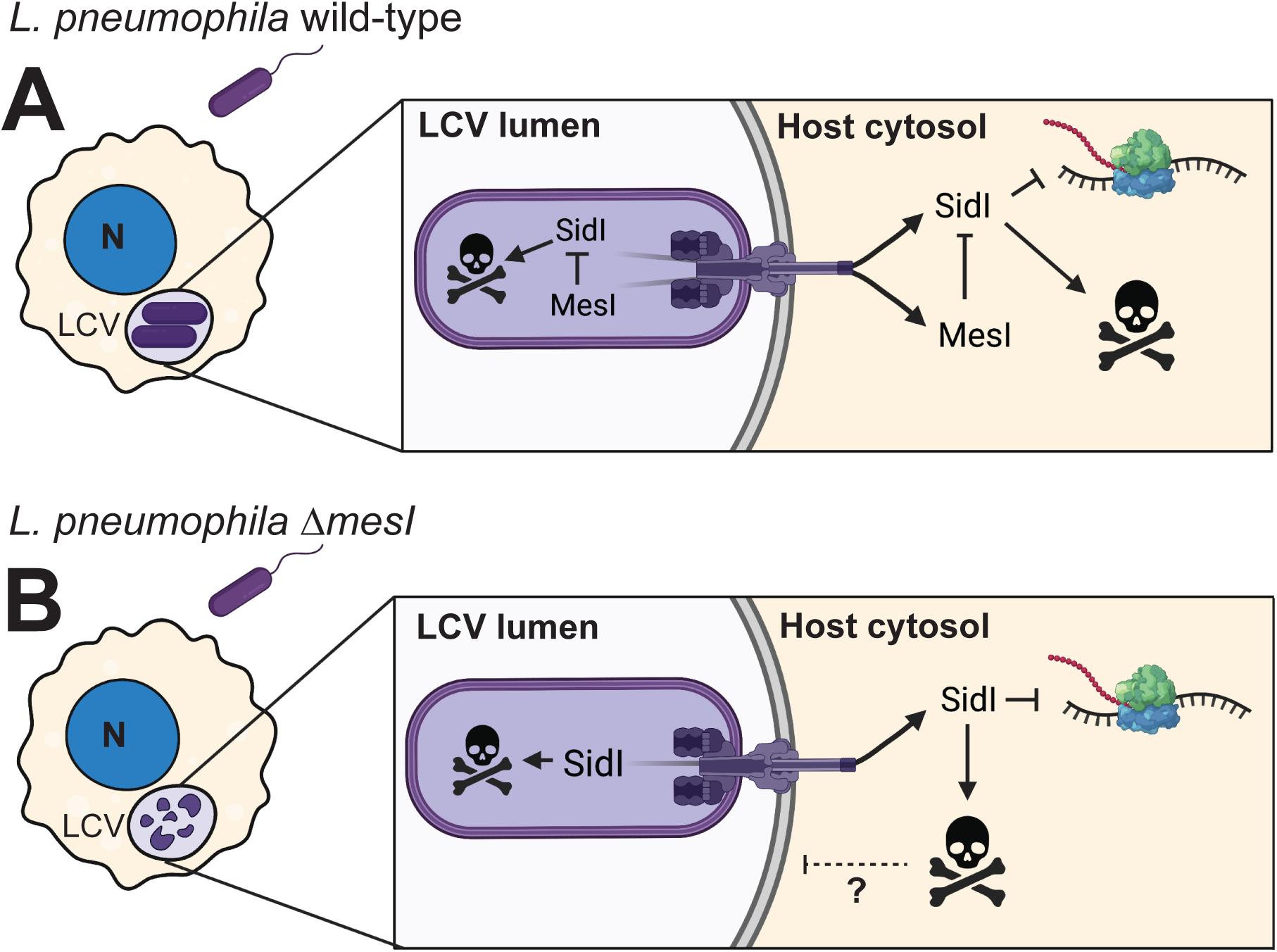
Model for the MesI regulatory mechanism. **(A)** MesI regulates SidI toxicity within host cells and the bacterium promotes replication of virulent *L. pneumophila* (purple) within host cells by regulating SidI within the bacterium and host cell. SidI inhibits eukaryotic protein synthesis and is toxic to both host cells and *L. pneumophila* in the absence of MesI. **(B)** MesI-deficient bacteria are degraded within host cells, which is likely a consequence of intrabacterial SidI toxicity. A role for SidI host cell toxicity in the *L. pneumophila* Δ*mesI* virulence defect is unclear (dashed line). Figure created with Biorender.com.

Interkingdom SidI toxicity suggests that its target is highly conserved between *L. pneumophila* and host cells. SidI is one of at least seven *L. pneumophila* effectors that subvert host protein synthesis (11, 31), one of the most conserved cellular processes across the kingdoms of life (32). Effector-mediated suppression of host mRNA translation is important for *L. pneumophila*’s acquisition of host-derived amino acids, which are essential for intracellular replication and the main driver of bacterial phase switching within host cells (23, 33). The mechanism by which SidI inhibits protein synthesis is unclear; however, SidI binds eEF1A and likely functions as a mannosyltransferase, so it is tempting to speculate that SidI blocks translation by modifying eEF1A (11, 12). However, the functional significance of eEF1A binding remains unclear since it is unaffected by MesI and insufficient for translation inhibition (11, 12). *L. pneumophila* has evolved strategies to prevent intrabacterial suppression of translation. For example, the effector Lgt1 blocks eukaryotic translation by glycosylating eukaryotic elongation factor (eEF)1A on Ser53, which is conserved in eukaryotes but not the bacterial eEF1A homolog, EF-Tu (34, 35). The mechanistic underpinnings of intrabacterial MesI activity is unclear; however, it is tempting to speculate that MesI may function in part to preserve protein synthesis within the bacterium.

Recent work has challenged the dogma that bacterial effectors function exclusively within host cells. Hardwidge and colleagues revealed that the pathogenic *E. coli* effector NleB and *Salmonella enterica* Typhimurium effector SseK1 function intrabacterially. NleB confers resistance to oxidative stress and SseK1 confers resistance to methylglyoxal and regulates UDP-GlcNAc biosynthesis (36–38). The pathogenic E. coli effector NleC can also function intrabacterially, but the endogenous, biologically relevant substrates are unknown (39). Our data suggest that intrabacterial MesI activity confers a fitness advantage on *L. pneumophila* by suppressing intrabacterial SidI activity. However, MesI is produced at later time points than SidI when *L. pneumophila* is grown in broth (19), suggesting that low levels of SidI activity within the bacterium may be beneficial at early stages of the *L. pneumophila* lifecycle. Indeed, there is emerging evidence that bacterial TA pairs can shape pathogen physiology by modulating post-transcriptional gene expression (40). However, this activity must be highly regulated by the antitoxin to prevent toxicity. Thus, it is tempting to speculate that SidI and MesI are an ancient TA pair that have been co-opted by the T4SS to subvert both host and pathogen physiology.

Our data challenge our initial assumption that MesI promotes *L. pneumophila* intracellular replication by preventing SidI-mediated host cell toxicity. We found no SidI-mediated differences in host cell death and observed that macrophages MesI-deficient bacteria had normal morphology. Interestingly, McCloskey *et al* recently proposed a model whereby MesI negatively regulates SidI translocation into host cells (19). MesI binds the extreme C-terminus of SidI, which encodes the canonical E-block Dot/Icm translocation signal (12, 19, 41). McCloskey *et al.* recently suggested that MesI negatively regulates SidI secretion from *L. pneumophila*. However, SidI secretion was not further attenuated when the relative abundance of MesI was increased (19). Furthermore, we did not detect any MesI-mediated differences in SidI translocation from *L. pneumophila* using an the established adenylate cyclase (CyaA) reporter (12). This discrepancy may be a consequence of differences in experimental conditions. Further investigation is required to define the impact of MesI on SidI translocation and the potential role of this activity in the MesI regulatory mechanism and *L. pneumophila* virulence.

Together, this study revealed a unique role for intrabacterial regulation of a translocated effector by its cognate metaeffector in *L. pneumophila* virulence. Our data, in the context of previously published studies (11, 12), suggest that SidI’s target is conserved between host and pathogen. Recent phylogenetic analysis suggests ancient and extensive co-evolution between the order *Legionellales,* which includes *Legionella* spp., and unicellular eukaryotes (42). Thus, metaeffectors may represent an ancient mechanism evolved by pathogenic bacteria for adaptation to eukaryotic hosts.

## Materials & Methods

### Bacterial strains, plasmids, primers, and culture conditions

*Escherichia coli* Top10 (cloning), DH5αλ*pir* (allelic exchange) and BL21 (DE3) (protein production) were maintained at 37°C in Luria-Bertani (LB) medium supplemented with appropriate antibiotics for plasmid maintenance [50 µg mL^-1^ kanamycin, 25 µg mL^-1^ chloramphenicol, or 100 µg mL^-1^ ampicillin]. Protein production in *E. coli* BL21 (DE3) was induced with 1 mM isopropyl-β-D-1-thiogalactopyranoside (IPTG).

All *Legionella pneumophila* strains in this study were derived from *L. pneumophila* Philadelphia-1 SRS43 (10) and cultured at 37°C on charcoal–N (2-acetamido)-2-aminoethanesulfonic acid (ACES)-buffered yeast extract (CYE) supplemented with L-cysteine and ferric pyrophosphate as described previously (43). Single colonies were isolated after 4 days of growth on CYE agar and used to generate 48 h heavy patches of bacteria. Where indicated, liquid cultures were grown from heavy-patched bacteria at 37°C in L-cysteine- and ferric pyrophosphate-supplemented ACES buffered yeast extract broth (AYE) (44). Media were supplemented with 10 µg mL-1 chloramphenicol (plasmid maintenance), 10 µg mL-1 kanamycin (allelic exchange), 1 mM IPTG (P*tac* gene expression), or 0.6-2% L-arabinose (w/v; paraBAD gene expression).

Plasmids and oligonucleotide primers used in this study are listed in **Table 1** and **Table 2**, respectively.

**Table 1.** Plasmids used in this study

| Name | Description | Resistance | Ref/Source |
| --- | --- | --- | --- |
| *Protein production* |  |  |  |
| pT7HMT | Production of His<sub>6</sub>-Myc fusions | Kan<sup>R</sup> | (47) |
| pT7HMT::*mesI* | Production of His<sub>6</sub>-Myc-MesI | Kan<sup>R</sup> | (12) |
| pT7HMT::*mesI*<sub>Δ10</sub> | Production of His<sub>6</sub>-Myc-MesI<sub>Δ10</sub> | Kan<sup>R</sup> | This study |
| pGEX6P1 | Production of GST | Amp<sup>R</sup> | GE healthcare |
| pGEX6P1::*sidI* | Production of GST-SidI | Amp<sup>R</sup> | (12) |
| *Allelic exchange* |  |  |  |
| pSR47s | Allelic exchange vector | Kan<sup>R</sup> | (48) |
| pSR47s::*paraBAD* | For chromosomal integration of *araBAD-3xflag* upstream of *sidI* | Kan<sup>R</sup> | This study |
| pSR47s::*ΔdotA* | For in-frame deletion of *dotA* | Kan<sup>R</sup> | This study |
| *Genetic complementation and cloning* |  |  |  |
| pJB1806 | *Legionella* expression vector |  | (45) |
| pJB1806::*sidI* (*psidI*) | *sidI* expression from *Ptac* | Cm<sup>R</sup> | This study |
| pJB1806::*sidI*<sub>R453P</sub> (*psidI*<sub>RP</sub>) | *sidI*<sub>R453P</sub> expression from *Ptac* | Cm<sup>R</sup> | This study |
| pSN85 | *Legionella* vector with in-frame N-terminal 3xflag epitope tag |  | (46) |
| pSN85::*sidI* | To generate 3xflag-*sidI* | Cm<sup>R</sup> | This study |
| pSN85::*mesI* (*pmesI*) | 3xflag-*mesI* expression from *Ptac* | Cm<sup>R</sup> | This study |
| pSN85::*mesI*<sub>Δ10</sub> (*pmesI*<sub>Δ10</sub>) | 3xflag-*mesI*<sub>Δ10</sub> expression from *Ptac* | Cm<sup>R</sup> | This study |

**Table 2.**
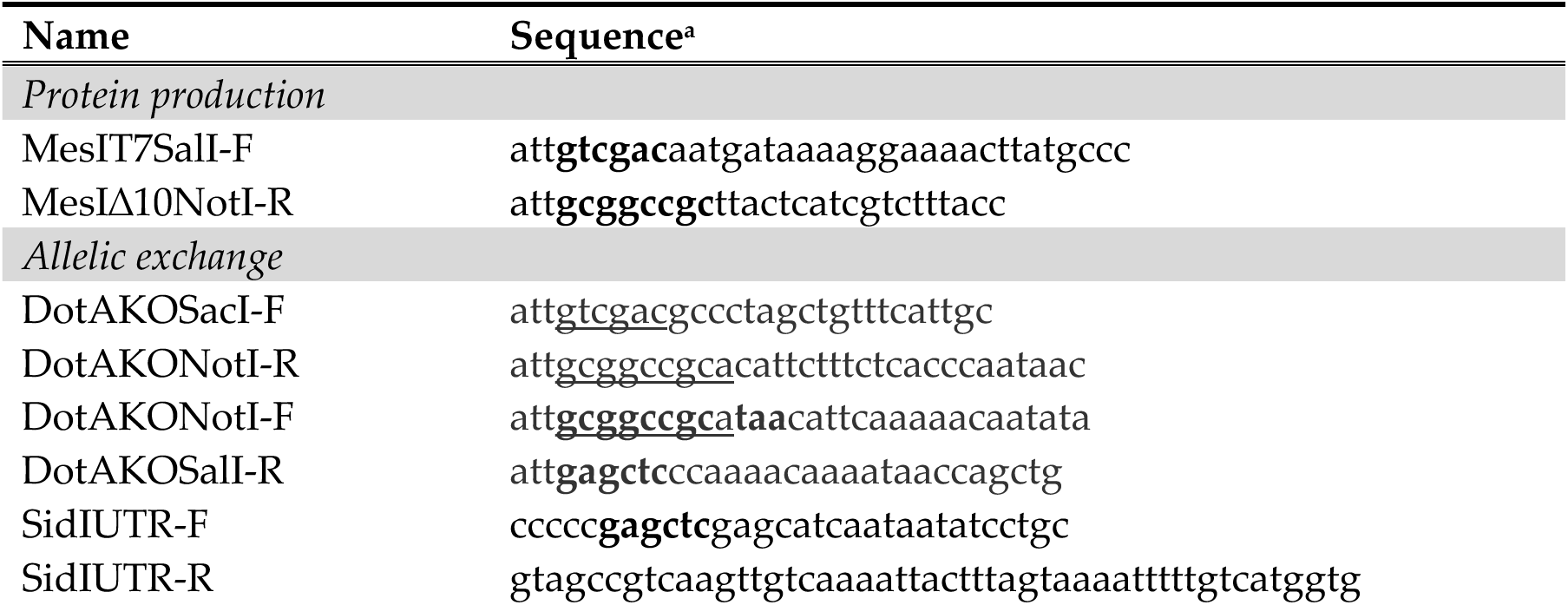
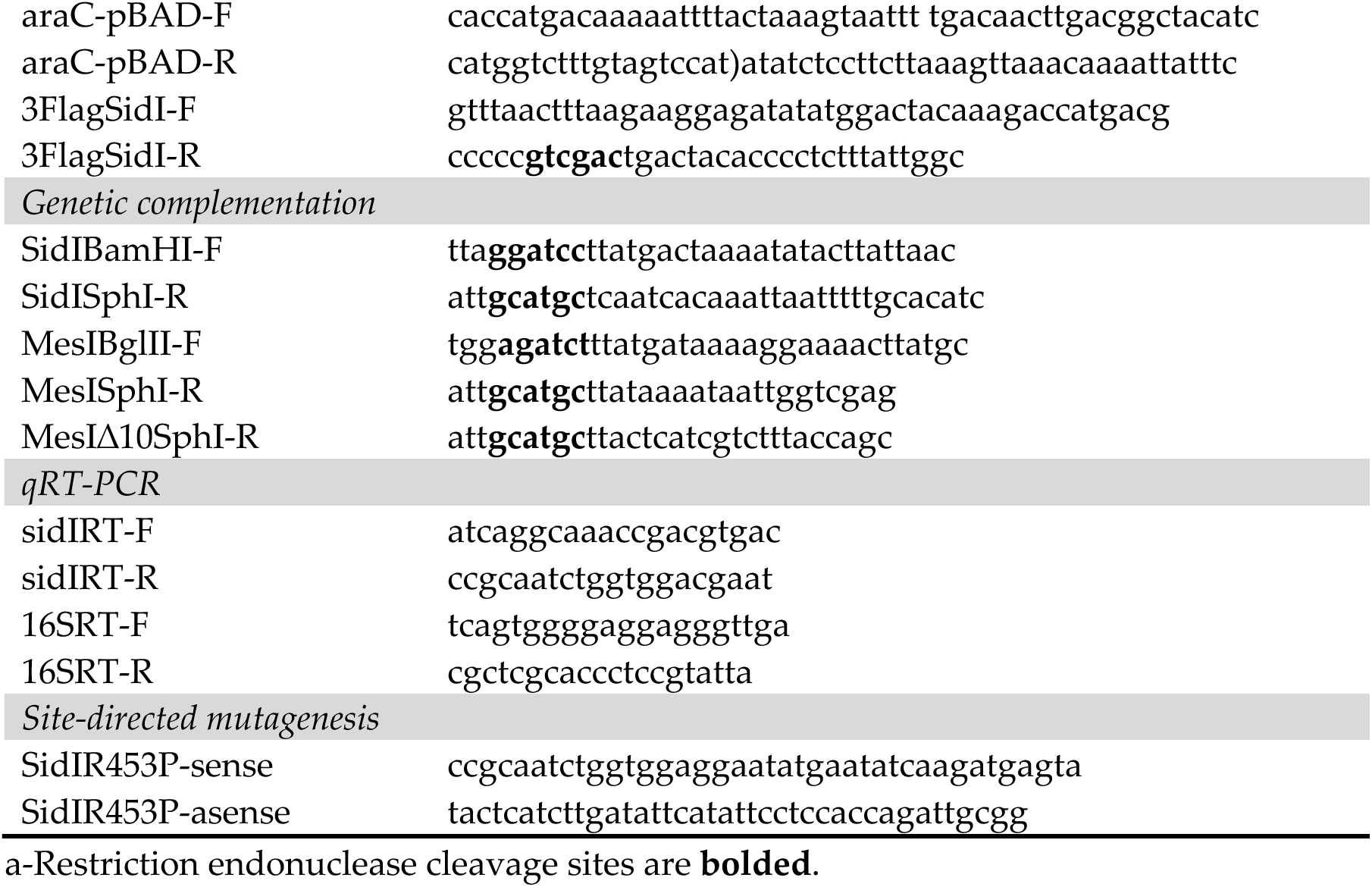
Oligonucleotide primers used in this study

### Molecular cloning and strain construction

*L. pneumophila* gDNA was isolated using the Illustra genomicPREP DNA spin kit (GE Healthcare) and used as a template for cloning *sidI* and *mesI* into the indicated plasmid vectors. *sidI* was amplified from *L. pneumophila* gDNA using SidIBamHIF/SidISphI-R, which includes 60 base-pairs (bp) downstream of the *sidI* open reading frame, and cloned as a BamHI/SphI fragment into pJB1806 (45) or downstream of an in-frame *3xflag* epitope tag in pSN85 (46) to generate pJB1806::*sidI* (p*sidI*) and pSN85::*sidI*. To generate p*sidI*_R453P_, p*sidI* was mutagenized using site-directed mutagenesis with primer pairs SidIR453P-sense/SidIR453Pasense (10). To generate p*mesI* complementation plasmids, *mesI* and *mesI*_Δ10_ open reading frames were amplified from *L. pneumophila* gDNA using MesIBglII-F/MesISphI-R or MesIBglII-F/MesIΔ10SphI-R primer pairs, respectively, and cloned into pSN85 in-frame with the *3xflag* epitope tag. Ligations were transformed into chemically competent *E. coli* Top10 and sequences confirmed by Sanger sequencing (Eton Biosciences). *L. pneumophila* complementation strains were generated by electroporation of plasmids into competent *L. pneumophila* strains using a BioRad Gene Pulser at 2.4 kV, 200 Ω, and 0.25 µF and plated on CYE supplemented with 10 µg mL^-1^ chloramphenicol.

For production of recombinant His_6_-Myc-MesIΔ10, *mesI*Δ10 was amplified from *L. pneumophila* gDNA using MesIT7SalI-F/MesIΔ10NotI-R and cloned as a SalI/NotI fragment into pT7HMT (47) downstream of an in-frame His_6_-Myc epitope tag. pT7HMT::*mesI* and pGEX::*sidI* were generated previously (**Table 2**) (12). Ligation reactions were transformed into chemically competent *E. coli* Top10 and sequences were confirmed by Sanger sequencing (Eton Biosciences).

The *dotA* open reading was deleted from the *L. pneumophila* chromosome using allelic exchange, as described (48). To generate a clean in-frame deletion of *dotA*, 5’ and 3’ regions flanking the *dotA* open reading frame (726 bp upstream and 1,012 bp downstream) were amplified using DotAKOSacI-F/DotAKONotI-R and DotAKONotI-F/DotAKOSalI-R to generate SacI/NotI and NotI/SalI fragments which were ligated into SacI/SalI digested pSR47s to generate pSR47s::Δ*dotA*, which was conjugated into *L. pneumophila* SRS43 for selection of double crossover events, as described. Sequences were confirmed by Sanger sequencing (Eton Biosciences) and the ΔdotA phenotype was confirmed by comparison the established SRS43 *dotA*::Tn strain (10).

For allelic exchange to insert the paraBAD promoter and a *3xflag* epitope tag upstream of *sidI* in the *L. pneumophila* chromosome, overlapping primers were used to generate a fusion construct consisting of 1,000 bp upstream of *sidI* (SidIUTR-F/SidIUTR-R; from *L. pneumophila* gDNA), araBAD promoter [araC-pBAD-F/araC-pBAD-R; from pMRBAD::z-CspGFP (a gift from Dr. Brian McNaughton; Addgene #40730) (49)], and *3xflag* fused to the first 1,200 nucleotides of *sidI* (3FLAG-SidI-F/3FLAG-SidI-R; from pSN85::*sidI*), which was ligated as a SacI/SalI fragment into pSR47s to generate pSR47S::araBAD. Plasmids were transformed into chemically competent *E. coli* DH5αλ*pir* and sequences were confirmed by Sanger sequencing (Eton Biosciences). To generate paraBAD strains, pSR47s::paraBAD was conjugated into *L. pneumophila* SRS43 wild type, Δ*mesI*, and *sidI*_R453P_Δ*mesI* (10) and selection of double crossover events was performed as described (48). Sucrose resistant, kanamycin sensitive colonies were screened by PCR to verify gene insertion.

### Mice

C57Bl/6 *Nlrc4*^-/-^ mice (a gift from Dr. Craig Roy) have been described (50). In-house colonies were maintained under specific pathogen-free conditions at Kansas State University. Bone marrow was harvested from 8- to 15-week-old mice as previously described (51). All experiments involving animals were approved by the Kansas State University Institutional Animal Care and Use Committee (IACUC-4501 and -4022) and performed in compliance with the Animal Welfare Act and National Institutes of Health guidelines.

### Cell culture

All mammalian cells were grown at 37°C/5% CO2. HEK293 FcγRII cells (a gift from Dr. Craig Roy) were cultured in Dulbecco’s Modified Eagle Medium (DMEM; Gibco) supplemented with 10% heat-inactivated fetal bovine serum (HIFBS; Biowest) for up to 25 passages. Primary mouse bone marrow-derived macrophages (BMDMs) were differentiated in RPMI supplemented with 20% HIFBS (Biowest) and 15% L-929 conditioned medium (Differentiation media) for 6 days prior to infection, as described (51). Differentiated BMDMs were seeded for infections in RMPI supplemented with 10%HIFBS and 7.5% L-929 conditioned medium (Seeding medium).

### Macrophage growth curves

BMDMs were seeded into 24-well tissue culture dishes at 2.5 x 10^5^ per well one day prior to infection. *L. pneumophila* were cultured on CYE agar and heavy patch-grown bacteria were used to infect BMDMs at a MOI of 1 in triplicates and CFU were enumerated as previously described (10, 15). Fold growth was calculated by normalizing CFU at 24 h, 48 h, and 72 h to internalized bacteria at 1 h post-infection. For genetic complementation, either 1 mM IPTG or 2% L-arabinose (w/v) were added to the media as indicated at the time of infection and maintained throughout.

### *In vitro* growth curves

*L. pneumophila* heavy patches were resuspended in fresh AYE broth and sub-cultured for consistent 600 nm optical density (OD_600_). Cultures were split into triplicate wells of a 96-well round-bottom plate and incubated at 37°C with continuous orbital shaking using an Agilent BioTek Epoch2 plate reader. OD_600_ from triplicate wells was read every two hours for 36 h. Plasmids were maintained with 10 µg mL^-1^ chloramphenicol and IPTG (1 mM) or 1% L-arabinose (w/v) were added to the media to induce gene expression, as indicated.

### Quantitative RT-PCR

*L. pneumophila* strains were grown for 1 or 6 h in AYE broth of incubation with and without arabinose (2%), and total RNA was purified with the Direct-zol RNA miniprep kit with TRI-reagent (Zymo Research) following manufacturer’s instructions. RNA samples were treated with DNase (Sigma) before reverse transcription with Superscript III (Invitrogen). Quantitative RT-PCR was performed using the Invitrogen SuperScript III Platinum SYBR Green One Step qRT-PCR kit. Transcript abundance was quantified using sidIRT-F/sidIRT-R (sidI) and 16SRT-F/16SRT-R (16S rRNA) primer pairs on a BioRad CFX96 Real-Time PCR machine. Fold expression (2^ΔΔCt) was calculated by normalizing *sidI* transcript abundance to 16S rRNA and standardizing values to wild-type *L. pneumophila*.

### Bacterial viability and LIVE/DEAD staining

*L. pneumophila* paraBAD strains were resuspended in AYE media from a heavy patch (48 h). Strains were grown for 24 h at 37°C in the presence or absence of 0.6% L-arabinose (w/v). Cell viability was quantified using a LIVE/DEAD™ *Bac-*Light™ Bacterial Viability kit according to manufacturer’s instructions. Bacteria were stained with 5 µM SYTO9 (live; green) and 10 µM propidium iodide (dead; red) (ThermoFisher) and incubated for 15 min at room temperature in the dark. To determine the baseline threshold for dead cells a negative control was used, where cells were treated with 90% ethanol for 1 h. After staining, cells were washed with sterile water and 5µL of resuspended cells were loaded on a glass slide with a coverslip. Images were acquired at the KSU Division of Biology Microscopy Facility using a Zeiss LSM5 Laser Scanning Confocal Microscope using a 100x oil-immersion objective. Percent dead bacteria was calculated by normalizing dead bacteria (red) to total bacteria using ImageJ software. At least 500 cells were evaluated for each strain and culture condition and the researcher performing the analysis was blinded to sample identity.

### Western blot

Protein was boiled in Laemmli Buffer and separated by SDS-PAGE and were transferred to polyvinylidene difluoride (PVDF; Fisher Scientific) membranes using a BioRad wet transfer apparatus. The membranes were incubated with blocking buffer [5% nonfat milk powder dissolved in Tris-buffered saline-0.1% Tween 20 (TBST)] for 30 min. Membranes were incubated with primary antibodies [mouse α-FLAG M2 (1:1000; Sigma); rabbit α-ICDH (1:1,000; Sigma); rabbit α-β-actin (Cell Signaling); rabbit α-Myc TAG mAb (71D10) (1:1,000; Cell Signaling)] at 1:1,000 in blocking buffer overnight at 4°C with rocking and subsequently with HRP-conjugated goat α-rabbit or α-mouse secondary antibodies at 1:5,000 (ThermoFisher) in blocking buffer for 2 h at room temperature with rocking. Membranes were washed, incubated with ECL reagent (GE Amersham), and imaged by chemiluminescence using an Azure Biosystems c300 Darkroom Replacer.

### Saponin solubilization assay

HEK293 FcγRII cells (52) were seeded in 10 cm poly-L-lysine-coated tissue culture dishes (4 x 10^6^) one day prior to infection. *L. pneumophila* strains were patched from fresh single colonies onto CYE agar supplemented with 10 µg mL^-1^ chloramphenicol and 1 mM IPTG to induce gene expression. Bacteria were opsonized with α-*L. pneumophila* antibody [1:1,000 (Invitrogen; PA17227)] in DMEM 10%HIFBS supplemented with 1 mM IPTG for 20 min at room temperature (RT) with rotation. Cells were infected with opsonized bacteria at a MOI of 50 in for 2 h in 2 mL DMEM 10%HIFBS supplemented with 1 mM IPTG. Cells were washed 3X with ice-cold PBS to remove extracellular bacteria and lysed for 10 min in 500 µL icecold Hank’s Balanced Salt Solution (HBSS) supplemented with 0.2% saponin and ProBlock™ Gold Mammalian Protease Inhibitor Cocktail (GoldBio). Lysates were treated with RNAse A (10 µg mL-1) and DNase I (10 µg mL-1), incubated at RT for 15 min, and centrifuged for 15 min at 17,000 r.c.f at 4°C. Saponin-soluble supernatants were filtered using a 0.22 µM syringe filter and transferred to a fresh 1.5 mL microcentrifuge tube. Saponin-insoluble pellets were resuspended in 50 µL TE buffer (10 mM Tris-HCl, 1 mM EDTA). Fifty microliters of supernatant and pellet fractions were diluted in 3X Laemmeli sample buffer, boiled for 10 min, and the whole sample was loaded into wells of a 1.5 mm 15% SDS-PAGE gel for Western blot.

### Affinity chromatography

Affinity chromatography from *E. coli* lysates was performed as described (12). Briefly, *E. coli* BL21 (DE3) harboring pT7HMT::*mesI*, pT7HMT::*mesI*_Δ10_, pGEX6P1::sidI, or pGEX6P1 were grown overnight with shaking at 37°C overnight and sub-cultured at 1:100 in LB media. Sub-cultures were grown for 3 h followed by induction with 1 mM IPTG and growth overnight at 16°C. Clarified lysates from bacteria producing GST-fusion proteins were incubated with pre-equilibrated glutathione magnetic agarose beads (Pierce) for 1h at 4°C with rotation. Beads were washed twice in washing buffer (125 mM Tris-Cl, 150 mM NaCl, 1mM DTT, 1 mM EDTA, pH 7.4) and incubated with clarified lysates from bacteria producing His_6_-Myc fusion proteins at 4°C for 1 h with rotation. Beads were washed twice in washing buffer transferred to a fresh 1.5 mL microcentrifuge tube and boiled in 25 µM 3X Laemmli Sample buffer. Proteins were separated by SDS-PAGE and visualized using Coomassie brilliant blue or Western Blot, as indicated.

### Cytotoxicity Assay

Cytotoxicity was evaluated by quantifying lactate dehydrogenase (LDH) activity in supernatants of *L. pneumophila*-infected BMDMs. BMDMs were seeded in 24-well tissue culture dishes at 2.5×10^5^ in Seeding medium one day prior to infection. Cells were infected with *L. pneumophila* strains at an MOI of 50 for 1 h, washed 3 times with sterile phosphate-buffered saline (PBS^-/-^), and incubated in seeding media for additional 5 h or 9 h. At the indicated times, plates were centrifuged at 250 r.c.f. and supernatants were transferred to a sterile non-tissue culture treated 96 well plate. For the 1 h timepoint, supernatants were collected from cells without washing. LDH was quantified using the CytoTox™ 96 Non-radioactive Cytotoxicity Assay (Promega) and according to manufacturer’s instructions. Absorbance at 490 nm was read on a BioTek Epoch2 microplate reader and percent cytotoxicity was calculated by normalizing absorbance values to cells treated with lysis buffer.

### Quantification of bacterial degradation within macrophages

BMDMs (1 x 10^5^) were seeded on poly-L-lysine-coated glass coverslips in 24-well tissue culture dishes one day prior to infection. BMDMs were infected in triplicate wells with the indicated *L. pneumophila* strains at an MOI of 30 for 1 h or 8 h. For the 8 h timepoint, extracellular bacteria were removed after 1 h by washing coverslips three times in PBS^-/-^ and incubated with fresh media for 7 h. Coverslips were fixed in 3.7% paraformaldehyde for 15 min and permeabilized with ice-cold methanol. Coverslips were stained with 1:1,000 rabbit α-*L. pneumophila* primary antibody (Invitrogen; PA17227) and 1:500 Alexa488-conjugated goat α-rabbit secondary antibody (ThermoFisher). Nuclei were stained with 1:2,000 Hoeshst (ThermoFisher) and coverslips were mounted on slides with ProLong™ Gold Antifade Mountant (Invitrogen). Degraded and intact bacteria were scored blind on a Leica DMiL LED inverted epifluorescence microscope (*n*=150 per strain and time point). Representative images were acquired at the KSU College of Veterinary Medicine Confocal Core using a Zeiss LSM 880 inverted confocal microscope and processed using Fiji ImageJ and Adobe Photoshop software.

### Statistical analysis

Statistical analysis was performed with GraphPad Prism 9 software using unpaired two-tailed *t*-test with a 95% confidence interval. Unless otherwise indicated, data are presented as mean ± standard deviation (s.d.) and statistical analysis was performed on triplicate biological replicates.

## Acknowledgements

We thank Dr. Craig Roy for providing *Nlrc4*^-/-^ mice and Dr. Philip Hardwidge for critical review of the manuscript and helpful discussions. This work was funded by NIH/NIGMS Center for Biomedical Research Excellence Research Project Grants (P20GM130448 and P20113117; to S.R.S.), an NIH/NIGMS Kansas-INBRE Postdoctoral Fellowship (P20GM103418; to D.C.), and start-up funds from Kansas State University.

## Supplemental Figures

**Figure S1.**
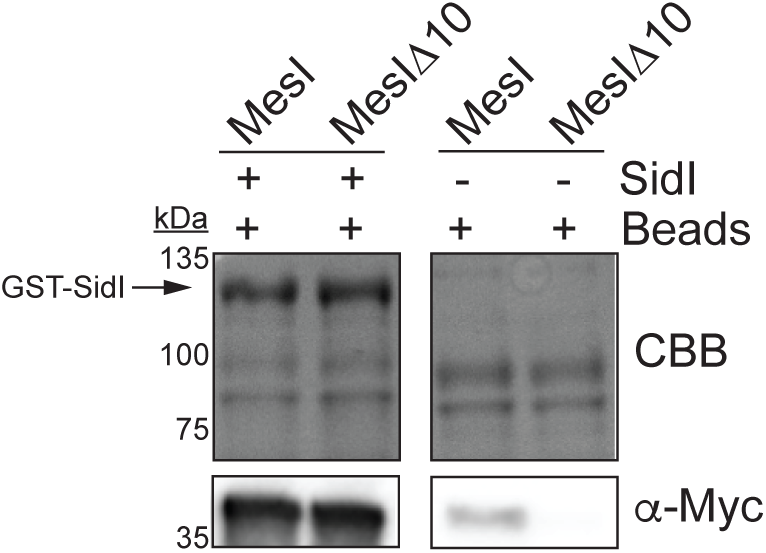
MesIΔ10 binds SidI. Glutathione agarose beads were incubated with lysates from *E. coli* producing GST-SidI or GST (beads) and subsequently incubated with lysates from *E. coli* producing His_6_-Myc-MesI or -MesIΔ10. Proteins on beads were separated by SDS-PAGE and visualized by Coomassie brilliant blue (CBB) staining or α-Myc Western blot, as indicated. Data are representative of three independent experiments.

**Figure S2.**
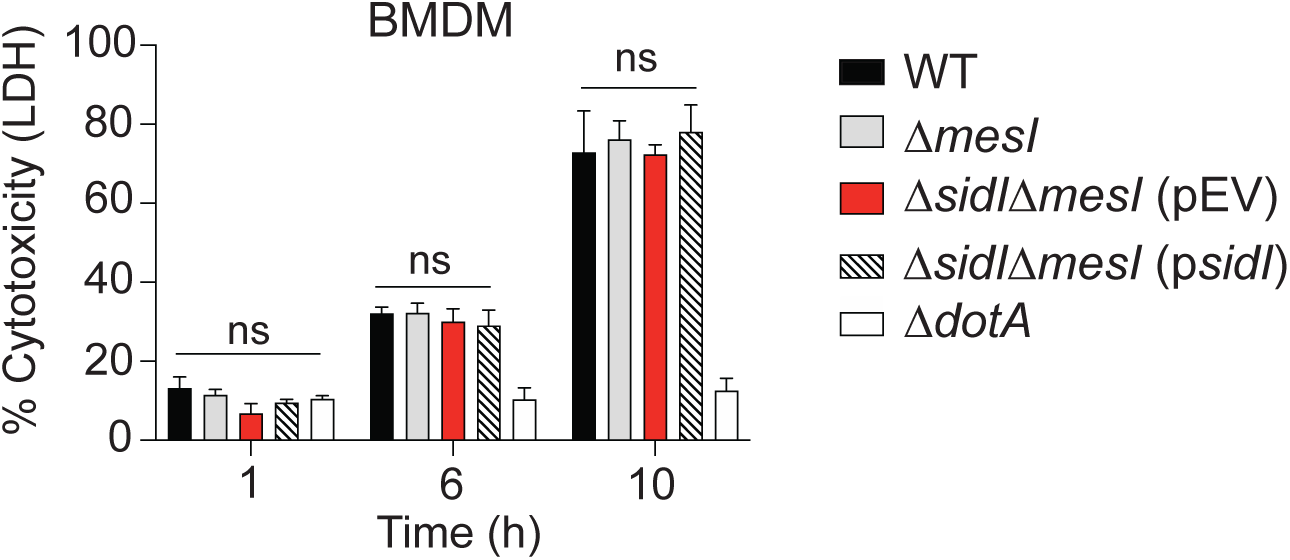
Host cell death is not increased by SidI-producing *L. pneumophila*. Lactate dehydrogenase activity in supernatants of *Nlrc4*^-/-^ BMDMs infected with the indicated *L. pneumophila* strains (MOI of 30) for 1, 6, or 10 h. Plasmid expression of *sidI* was induced with 1 mM IPTG. Data are representative of three independent experiments and statistical analysis was performed using Students’ *t*-test (ns, not significant).

## Notes

### Competing Interest Statement

The authors have declared no competing interest.

